# Sequencing coverage analysis for combinatorial DNA-based storage systems

**DOI:** 10.1101/2024.01.10.574966

**Authors:** Inbal Preuss, Ben Galili, Zohar Yakhini, Leon Anavy

## Abstract

This study introduces a novel model for analyzing and determining the required sequencing coverage in DNA-based data storage, focusing on combinatorial DNA encoding. We explore the application of the coupon collector model for combinatorial-letter reconstruction, post-sequencing, which ensure efficient data retrieval and error reduction. We use a Markov Chain model to compute the probability of error-free reconstruction. We develop theoretical bounds on the decoding probability and use empirical simulations to validate these bounds. The work contributes to the understanding of sequencing coverage in DNA-based data storage, offering insights into decoding complexity, error correction, and sequence reconstruction. We provide a Python package that takes the code design and other message parameters as input, and then computes the required read coverage to guarantee reconstruction at a given desired confidence.

## I. INTRODUCTION

**T**HE growing volume of the world’s digital data and the limitations of existing storage technologies motivate the need for new and innovative storage solutions [1].

DNA-based data storage (or DNA-based storage) emerges as a viable solution, offering unmatched density and durability. This novel approach involves the synthesis, storage, and sequencing of DNA molecules to encode, store and retrieve information. However, challenges such as short, error-prone strands and limitations in current synthesis technologies still remain [2] [3] [4] [5] [6] [7] [8].

While DNA-based storage stands as a promising technology, and the cost of DNA sequencing has been decreasing, it remains significantly more expensive than reading from established archival storage solutions [9] [10] [11] [12]. In the context of DNA sequencing costs and throughput, recent work [13] defined the DNA coverage depth problem, which considers the expected sample size, to guarantee successful decoding of the information. A related concept was suggested by Chandak et al. [14], who explored the balance of writing and reading costs in DNA-based data storage, studying the LDPC-based coding schemes.

Combinatorial DNA encoding is a recently introduced encoding scheme, which uses a set of clearly distinguishable DNA shortmers to construct large combinatorial alphabets, where each letter is encoded by a subset of shortmers [15]. The nature of these combinatorial alphabets minimizes mix-up errors, while also ensuring the robustness of the system. combinatorial shortmer encoding is an extension of other composite coding schemes, such as [2] [16] [17].

This work presents the first model for analyzing the sequencing coverage depth problem under combinatorial DNA encoding. In this work, we define and study a new model to compute the required coverage depth. While the model presented in [13] assumes 1D encoding applied on the strands, our model considers 2D (inner-outer) MDS codes. Each sequence is encoded using the inner-code, to protect against symbol errors, while the outer-code adds redundancy to a block of sequences, protecting against sequence-level errors. This allows for a more thorough and detailed analysis of the required sequencing coverage when using the inner-outer code approach, a widely used coding technique in DNA-based storage [18] [19] [3] [2].

We first address the question of reconstructing a single combinatorial letter by utilizing a reduction of this problem to the well-known coupon collector’s problem. This provides a framework for determining the required number of reads to ensure that at least one copy of every member shortmer in the combinatorial letter is observed [13] [20] [21] [22] [23]. For this purpose, we present a Markov Chain approach to calculate decoding probabilities and provide computer code. We also generalize this model by considering a threshold for the minimum number of copies of each shortmer required for letter reconstruction.

Next, we analyze the decoding probabilities of full-length combinatorial sequences that constitute a single complete message encoded using combinatorial DNA. We provide bounds on the decoding probability given the number of analyzed reads, and present an operational algorithm for determining the required coverage of reads. We explore our coverage depth model on different design parameters and compare the results to simulation experiments of combinatorial DNA reading.

Lastly, we provide computer code implementing our coverage model that, given a sequence and message design, outputs the read coverage required for recovering the data with a user defined confidence level. This work combines theoretical progress represented by studying the coverage depth problem for combinatorial DNA-based storage, and also the practical aspect supporting the design and implementation of such systems.

## II. Results

The decoding complexity is analyzed here by breaking the process down into its basic components. First, the decoding probability of a single combinatorial letter is analyzed, considering various design parameters and decoding approaches. Next, this paper addresses the decoding of a single combinatorial letter, while considering the use of error correction codes with varying redundancy levels. Finally, the decoding of a complete combinatorial DNA message is analyzed, considering a general 2D error correction MDS code (i.e., a code that protects against sequence dropouts as well as errors on each sequence).

### A. Reconstruction of a single combinatorial letter

Let Ω be a set of valid k-mers used for a combinatorial DNA-based data storage system. Consider a binomial combinatorial alphabet Σ with 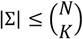 letters where each letter *σ* ∈ Σ consists of a subset of size *K* of k-mers from Ω. This subset is referred to as the member k-mers of *σ*. Let *R* be the number of analyzed reads of a given combinatorial letter. We define a decoding algorithm in which we accumulate reads until we observe at least *t* copies of *K* unique k-mers from Ω. These *K* k-mers are refrred to as the inferred member k-mers, and are used to reconstruct a combinatorial letter *σ′* (See Algorithm 1).

#### Algorithm 1: Decoding Algorithm for a Single Combination Letter

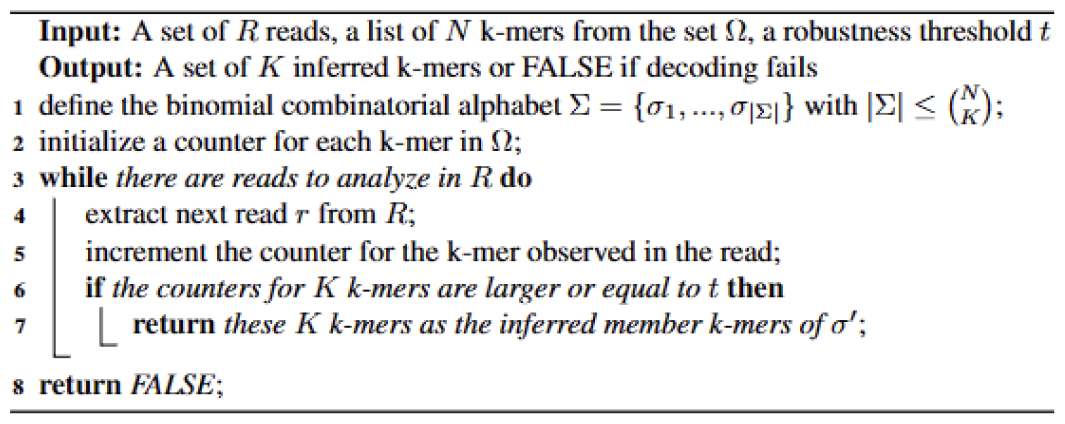

To analyze the probability of decoding a single combinatorial letter, we first assume that each read uniformly draws one of the *K* member k-mers. Let *T*(*K, t*) be a random variable representing the number of reads analyzed until the decoding algorithm successfully stops. Let *π*(*K, t*)(*R*) be the probability of stopping with a successful inference after at most *R*_*single*_ reads.

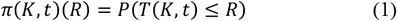

For *t* = 1 the random variable *T*(*K, t*) represents the classical coupon collector model [24] and we get (See Appendix B:

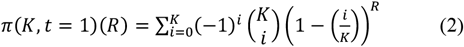

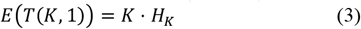

where 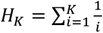 is the K^th^ harmonic number.

For *t* > 1 we can obtain [23]:

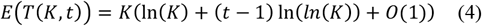

To calculate *π*(*K, t*)(*R*) for *t* > 1, we use a Markov Chain (MC) formulation. Each state in the MC represents the status of the member k-mers in *σ*, in terms of the number of times each has been seen. Specifically, a state is represented by a vector:

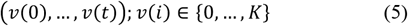

for 0 ≤ *j* < *t, v*(*j*) indicates the number of member k-mers seen exactly *j* times, while *v*(*t*) indicates the number of member k-mers seen *t* times or more.

Clearly, this vector satisfies:

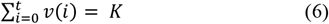

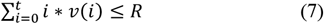

and when *v*(*t*) = 0 the inequality in (7) hold as equality 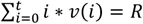. We also note that since there are *t* + 1 values in the vector (*v*(0), *v*(1), …, *v*(*t*)), there are a total of 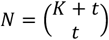 possible solutions to the equation, representing *N* states.

For example, considering *K* = 10 member k-mers and a threshold *t* = 2. The following states can be defined:

(10,0,0): All 10 k-mers have not been seen yet. This is the case before we start analyzing the reads.

(8,2,0): After analyzing two reads, two unique k-mers have been observed exactly once while the remaining 8 k-mers have not been observed yet.

(7,2,1): After analyzing at least four reads, two unique k-mers have been observed exactly once, one k-mer has been observed 2 times or more and the remaining 7 k-mers have not been observed yet.

We define the following transition matrix *A* where each transition is defined by the observation of one read.

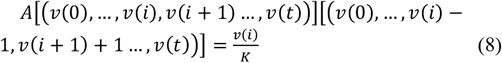

This represents observing one of the *v*(*i*) k-mers that were observed *i* < *t* times.

And,

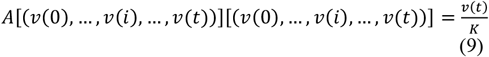

This represents observing one of the *v*(*t*) k-mers that were observed at least *t* times.

For example, the first two transitions are:

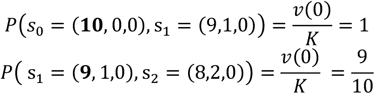

This happens when one out of the 9 yet unseen k-mers is drawn.

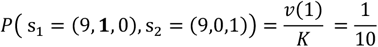

This happens only when the first observed k-mer is observed again.

To get *π*(*K, t*)(*R*) we set the initial state to be

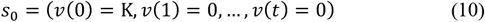

That is *P*_0_ = (*P*(*s*_0_) = 1,0, …,0) is the state distribution vector.

We derive the distribution vector over the states after *R* steps,

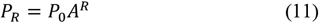

Let *s*_*f*_ = (0,0, …, *v*(*t*) = *K*) be the desired state in which all *K* k-mers have been observed at least *t* times.

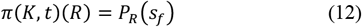

Fig. 1 and Appendix A. demonstrate the state distribution vector for several values of *R* using *K* = 5 member k-mers and a threshold of *t* = 1. Clearly, after analyzing the first read, a single k-mer is observed once while the other four have not been observed yet. With *R* = 5, the probability of having seen all unique coupons reached 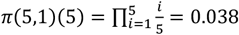 . At *R* = 15, this probability significantly increased to *π*(5,1)(15) = 0.829. Finally, at *R* = 30, the probability of observing all coupons was *π*(5,1)(30) = 0.994.

**Fig. 1.**
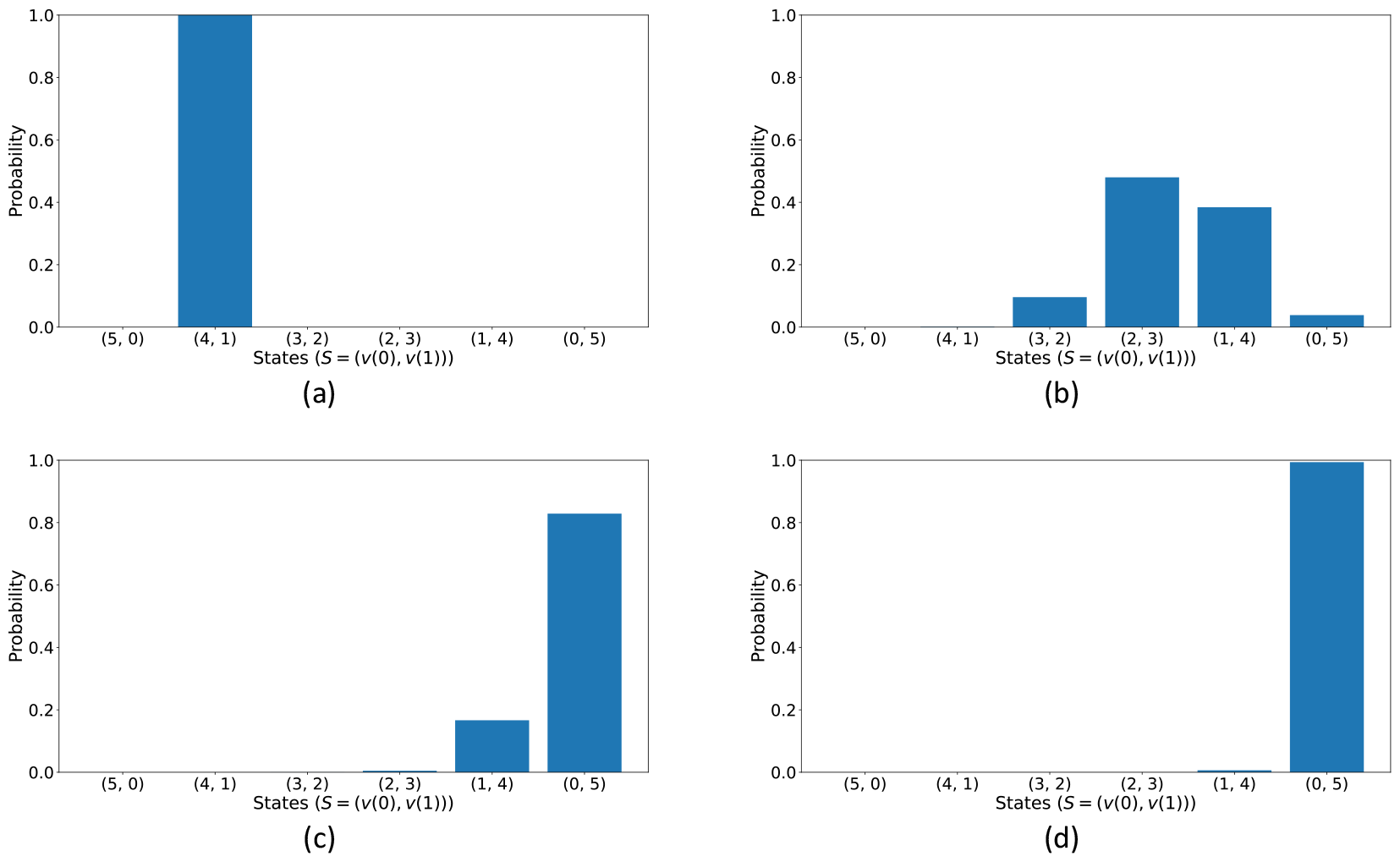
Evolution of probability in the coupon collector model. (a) The probability distribution across the 6 states (X-axis) after observing *R* = 1 reads. (b-d) Similar to (a) with *R* = 5, 15, 30 respectivley. Calculated for *K* = 5, *t* = 1, and no errors, *ϵ* = 0.

This algorithm ignores possible synthesis and sequencing error as it assumes that all observed k-mers come from the set of *K* valid k-mers. Introducing an error probability *ϵ* of observing an invalid k-mer requires a modified transition matrix *B*:

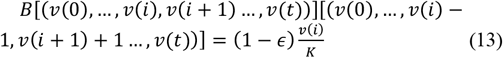

This represents observing one of the *v*(*i*) (valid) member k-mers that were observed *i* < *t* times.

And,

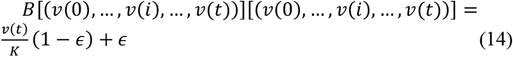

This represents observing one of the *v*(*t*) k-mers that were observed at least *t* times, or observing an invalid k-mer.

For example, the first two transitions are:

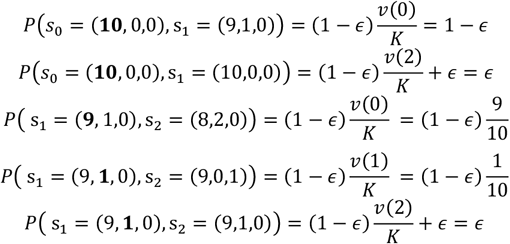

Fig. 2 depicts the decoding probabilities for varying number of analyzed reads using different values for the threshold *t*. The calculated probabilities are compared to a simulation experiment. As expected, as *t* increases, more reads are required to reconstruct a combinatorial letter. Notably, when *R* reaches 100 or more, the probability effectively becomes 1, indicating full data recovery. This represents the balance between threshold level required for achieving precise combinatorial reconstruction and the read depth complexity.

**Fig. 2.**
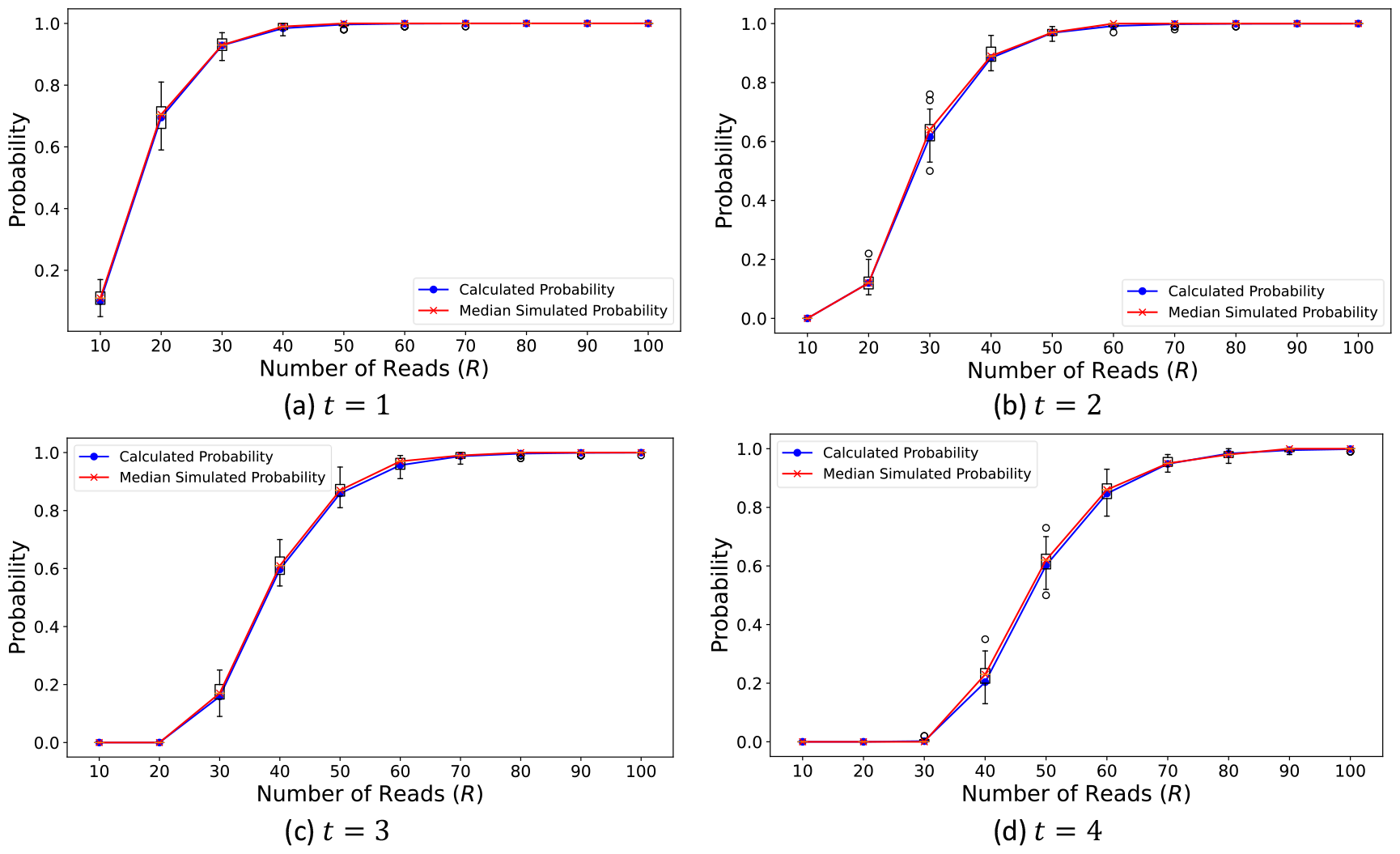
Decoding probability for varying number of analyzed reads (*R*) for different thresholds (*t*). Each subplot corresponds to a different threshold value (*t*). The analyses were conducted for *K* = 7 and *ϵ* = 0.01. (a) results for *t* = 1, the blue line corresponds to the calculated probability based on the MC model while the red line represents the median of 50 simulation runs, where each simulation calculates the success rate of 100 uniform drawing of *R* reads across *K* member k-mers. The simulation results are also presented as boxplots. (b-d). Like (a) with *t* = 2, 3 and 4 respectivley.

Note that throughout this section, we ignored the possibility of an error that results in k-mer mix-up (i.e., the output of the decoding algorithm is different from the original combinatorial letter, *σ*^*′*^ ≠ *σ*). This is due to the assumptions that the design parameters render this error type very unlikely. We furtherdiscuss this issue in Section IV. Discussion.

### B. Reconstruction of a combinatorial sequence

Let *s* = *σ*^(1)^*σ*^(2)^ … *σ*^(*m*)^ be a sequence of length *m* over the same binomial alphabet defined in the previous section. Assuming the use of a proper MDS error correction code, we say that decoding only *b* ≤ *m* letters is sufficient for decoding the complete sequence. Let *R* be the number of analyzed reads, fixing *K* and *t*, we denote *π*(*K, t*)(*R*) as *π*(*R*). Let *W* be a random variable representing the number of letters in *s* that were decoded. Assuming independence between the letters in *s* we get

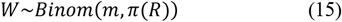

We are interested in the probability of decoding the sequence *s, P*_*single*_ :

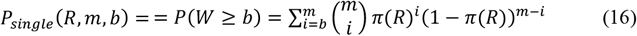

We can approximate this probability using the normal estimation (based on Central Limit Theorem).

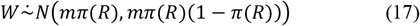

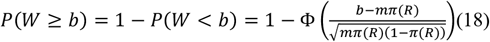

Where Φ is the CDF of the standard normal distribution.

Fig. 3 presents the decoding probabilities of a combinatorial sequence with length *m* = 100, examining how the number of analyzed reads (*R*) affects the accuracy of sequence reconstruction across various redundancy levels (*b* = 100, 95, 90, 85) keeping other parameters constant (*K* = 7, *t* = 4). We observe that the probability of successful reconstruction varies significantly with different redundancy levels. Notably, higher redundancy levels (lower *b* values) enable accurate reconstruction using fewer reads. These results also align with the results obtained from the Normal Approximation (Not shown). The results demonstrate the role of sequence level redundancy in affecting the likelihood of accurate reconstruction, making it an important tunable parameter in the overall design.

**Fig. 3.**
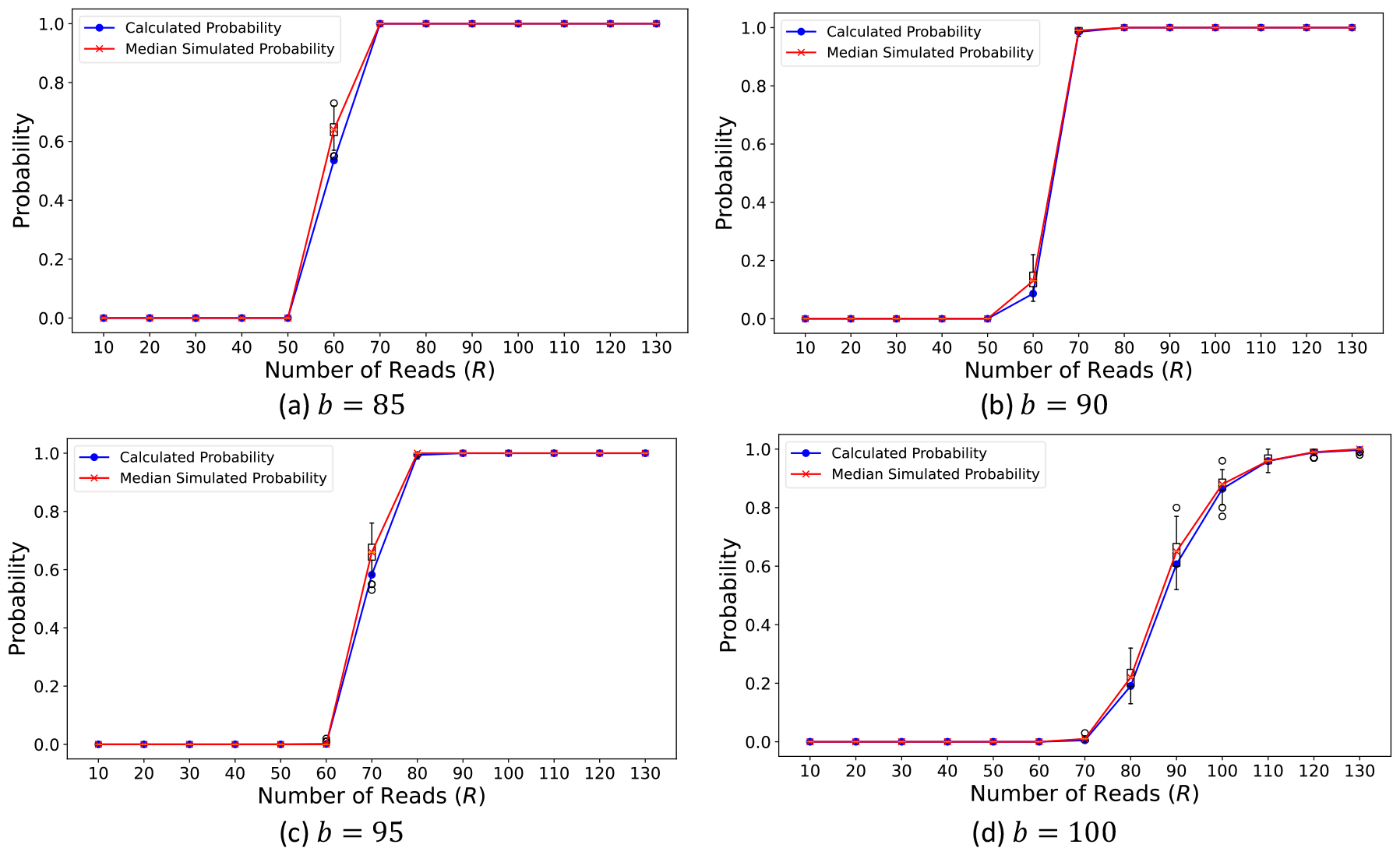
Decoding probability of a complete combinatorial sequence with varying redundancy levels. Results shown for a sequence of length *m* = 100, with *K* = 7, and requiring *t* = 4. (a) Calculated decoding probability (blue line) as a function of the number of analyzed reads for redundancy level of *b* = 85. Median results from 50 simulation runs are presents (red line) with boxplots representing the distribution of the simulation results. Each simulation run represents 100 uniformly drawn sets of *R* reads, each comprising *m* letters drawn from *K* = 7 member k-mers. (b-d) Like (a) with *b* = 90, 95, and 100, respectively. All analyses incorporate an error rate of *ϵ* = 0.01.

#### C. Reconstructing a complete combinatorial message

Let 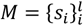 be a complete combinatorial message encoded using a binomial alphabet like in the previous sections. The message is encoded using *l* combinatorial sequences and, assuming proper MDS error correction code, *a* ≤ *l* of which are sufficient for the decoding of *M*.

Let *R*_*all*_ be the total number of analyzed reads over all sequences. We are interested in the probability of decoding at least *a* of *l* sequence using *R*_*all*_ reads, *P*_*all*_ (*R*_*all*_, *l, a*).

Fig. 4 presents an overview of the decoding process and the analysis steps for a complete combinatorial message. First, the *R*_*all*_ reads are distributed between the *l* sequences, using, for example, the barcodes. Then, the decoding probability of each of the *l* sequences is determined using the derivation from the previous section. The decoding probability of a single letter is analyzed using the coupon collector’s model. We now formally define each of these steps and analyze the decoding probability *P*_*all*_ (*R*_*all*_, *l, a*) or simply *P*_*all*_ .

**Fig. 4.**
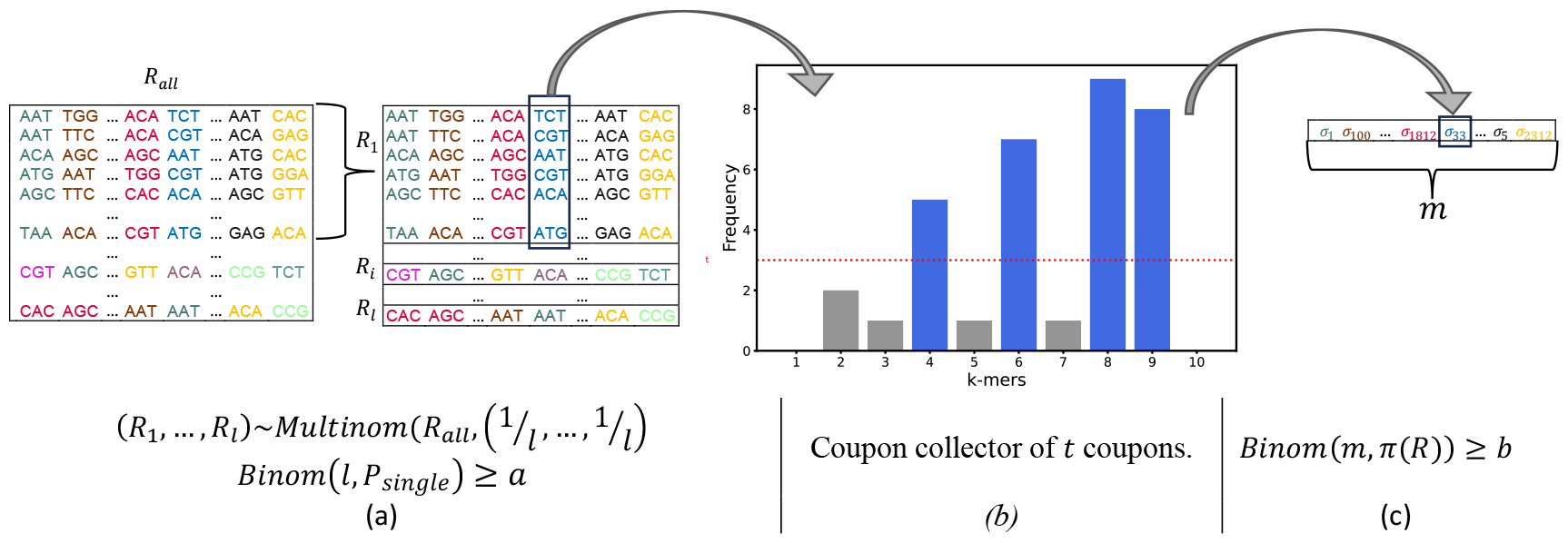
Reconstructing a complete combinatorial message. (a) *R*_*all*_ reads are distributed between *l* sequences and at least *a* sequences need to be decoded (b) The decoding probability of each of the letters is analyzed using the coupon collector’s model (Blue bins indicate the members k-mer) (c) Each sequence requires *b* of the *m* combinatorial letters to be decoded.

The distribution of the *R*_*all*_ reads across the *l* sequences is modeled using a multinomial distribution

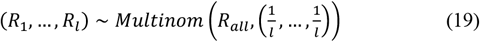

Given a specific distribution of the *R* reads (*r*_1_, …, *r*_*l*_), to successfully decode the message we need to decode at least *a* of the sequences.

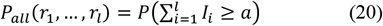

Where *I*_*i*_ is an indicator of decoding sequence *s*_*i*_ using *r*_*i*_ reads.

Using the law of total probability and setting *P*(*R*_1_ = *r*_1_, …, *R*_*l*_ = *r*_*l*_) = *P*(*r*_1_, …, *r*_*l*_) :

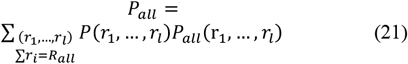

Calculating *P*_all_ directly becomes infeasible even for small values of *R*_*all*_, *l* and *a*. We therefore bound this probability.

First, we note that since for every sequence *s*_*i*_ we have

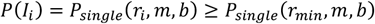

Where 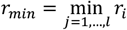 and 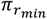 is obtained by using *r*_*min*_ in the coupon collector’s model. If we plug this back to (20) we can define a new binomial random variable *X* that represents the number of sequences decoded:

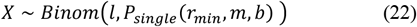

And,

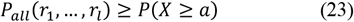

Yielding a lower bound on *P*_*all*_ (*R*_*all*_, *l, a*)

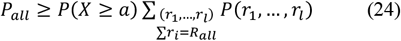

In the multinomial distribution for (*R*_1_, …, *R*_*l*_), many possible read distributions are very unlikely. We can further bound *P*_*all*_ by setting a constant value *ρ* and only considering read distributions for which 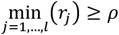 . Let *X*_*ρ*_ be a random variable representing the number of sequences decoded when the decoding probability of each sequence is calculated using *ρ* reads. That is, *X*_*ρ*_*∼Binom* (*l, P*_*single*_ (*ρ, m, b*)).

We therefore have

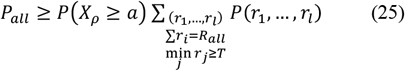

Given a small *δ* > 0, we check whether *R*_*all*_ reads are sufficient to decode the message with 1 − *δ* confidence level.

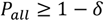

This can be achieved by choosing *ρ* such that

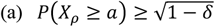

And,

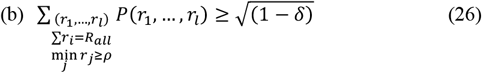

Since *X*_*ρ*_ has a binomial distribution, we can find *ρ* for which condition (a) holds. For condition (b), we use Sanov’s Theorem on the multinomial distribution as follows. For more on Sanov’s Theorem and the behavior of multinomials, see [25].

Sanov’s theorem bounds the probability that the distribution of the reads into barcodes significantly deviates from the expected uniform (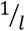 for each) distribution, particularly where at least one sequence gets fewer than *ρ* reads. Fig. 5 demonstrates this using a simulation of 100,000 instances each drawn from the multinomial distribution with 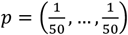 and *n* = 4500 or *n* = 5000. The plots show the distribution of the minimal values obtained. Clearly, increasing *R*_*all*_ reduces the probability of the minimal value to be below a fixed threshold *ρ*. Decreasing the threshold *ρ* yields a similar effect.

**Fig. 5.**
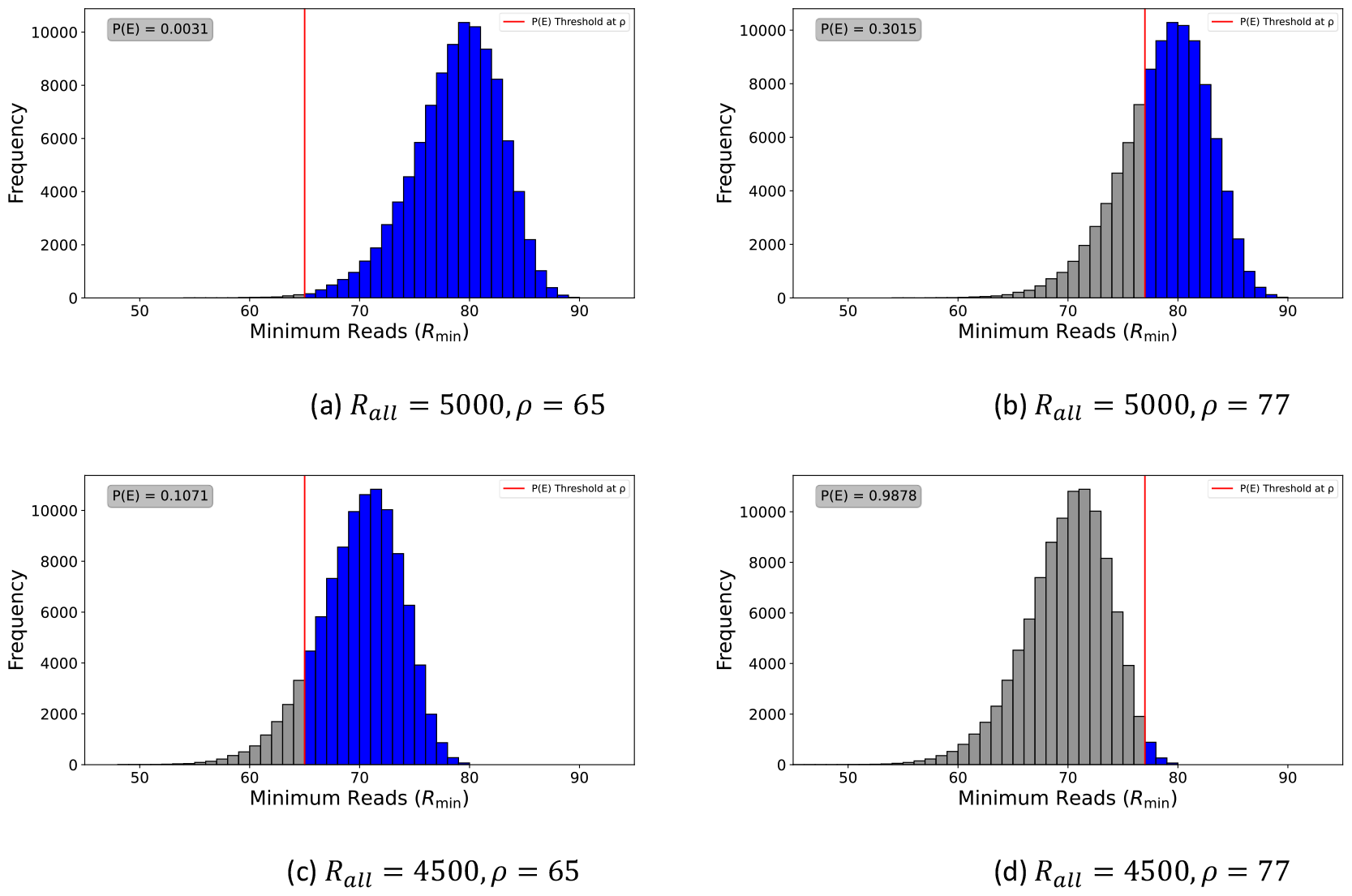
The minimum value of a multinomial distribution 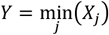 where 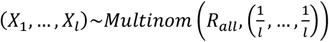. (a) A histogram of the values of *Y* attained in 100,000 instances with *l* = 50 and *R*_*all*_ = 5000. The red line represents *ρ* = 65. The gray box show the probability *P*(*E*(*ρ*)) = *P*(*Y* < *ρ*). (b-d) Like (a) for (*R*_*all*_, *ρ*) = (5000, 77), (4500, 65), (4500, 77).

Let 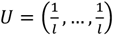 be the expected uniform distribution equivalent to the expected read distribution for (*R*_1_, …, *R*_*l*_).

Let *E*(*ρ*) be the set of probability vectors equivalent to read distributions (*r*_1_, …, *r*_*l*_) for which 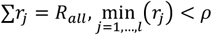 :

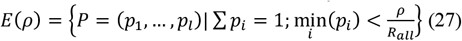

We define *ζ*(*ρ*) = *min*_*P*∈*E*_(_*T*_)*D*(*P* ∥ *U*) where *D*(*P*||*U*) is the Kullback-Leibler divergence:

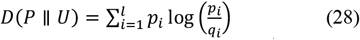

Let 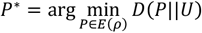 the closest element to *Q* in *E*(*ρ*) in terms of the KL-divergence. That is *ζ*(*ρ*) = *D*(*P*^*\**^ ∥ *U*)

Next we show that *P*^*\**^ is the distribution of reads in which *ρ* − 1 reads are assigned to one sequence and the remaining *R*_*all*_ − *ρ* + 1 reads are uniformly distributed over the remaining *l* − 1 sequences.

Lemma:

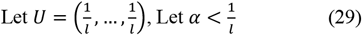

Let 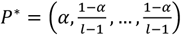, then

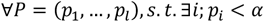

We have

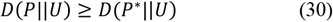

The proof for this lemma is found in the appendix C. For intuition, this is simply the result of the symmetric nature of the KL-divergence function and of *Q*.

Sanov’s Theorem [25] provides a bound on the probability of observing any distribution within *E*(*ρ*).

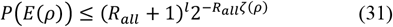

Where,

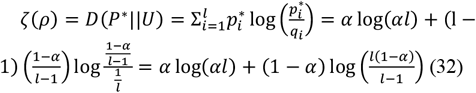

This bound implies that the likelihood of observing a significantly non-uniform distribution of reads decreases exponentially as the total number of reads *R*_*all*_ increases.

We recall that

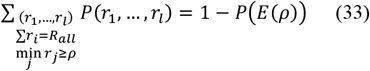

And so we get

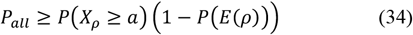

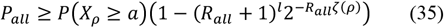

This gives us an operational algorithm for checking if *R*_*all*_ reads are sufficient to ensure successful decoding with confidence 1 − *δ*, as specified in Algorithm 2.

##### Algorithm 2: Finding required sequencing depth *R*_all_ for a complete message

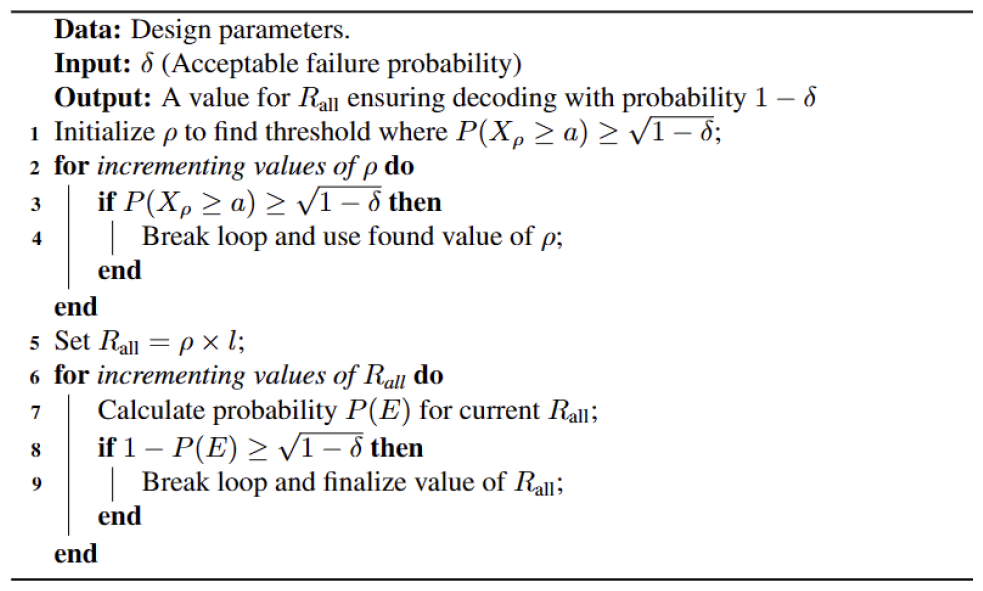

Fig. 6a demonstrates the approach by presenting the probability of successful message decoding *P*(*X*_*ρ*_ ≥ *a*) and the probability of considering “enough” of the read distribution (1 − *P*(*E*)) for a fixed number of overall reads *R*_*all*_ = 3000 as a function of the threshold *ρ*. Clearly, *P*(*X*_*ρ*_ > *a*) increases as *ρ* increases since each sequence *s*_*i*_ is decoded using more reads. On the other hand, as was demonstrated in Fig. 5. increasing *ρ* decreases 1 − *P*(*E*(*ρ*)) since less read distributions with 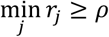 are expected.

**Fig. 6.**
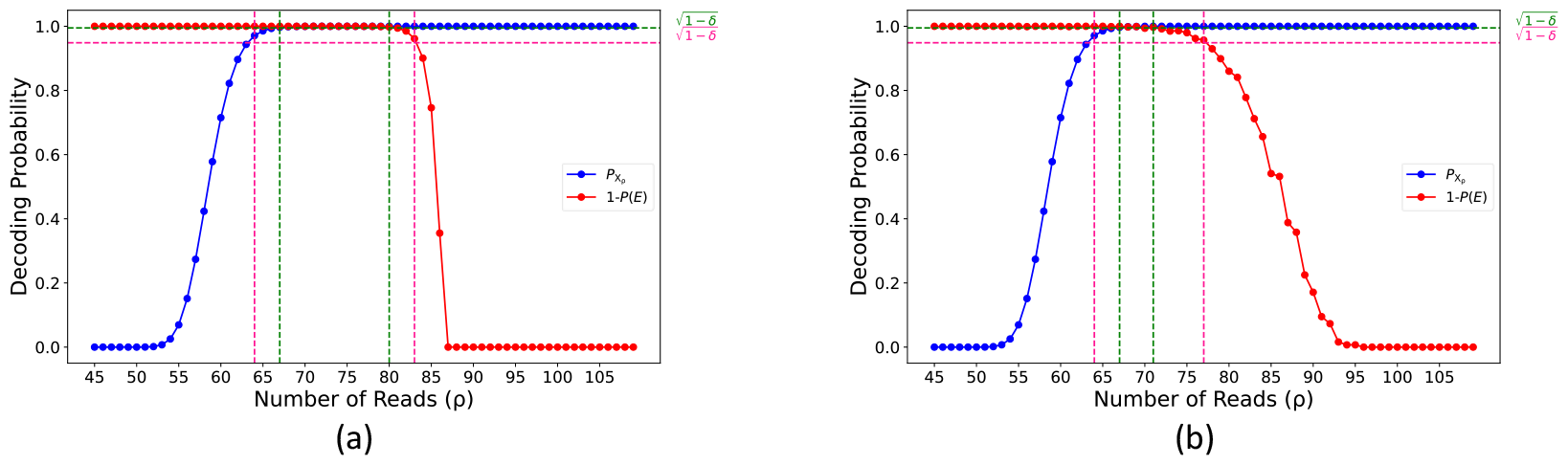
Bounding the decoding probability. (a) Overall decoding probability *P*(*X*_*ρ*_ ≥ *a*) (blue line) and the Sanov bound on the probability of obtaining a read distribution across the sequences with 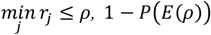 (red line) as functions of the threshold *T* for a fixed number of analyzed reads *R*_*all*_ = 3000. The threshold 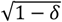 on the probability is marked with dotted lines for *δ* = 0.1 (pink dotted lines) and *δ* = 0.2 (green dotted lines), Setting *ρ* to any value between these lines ensures decoding with 1 − *δ* confidence. All values are calculated for *K* = 7, *t* = 4, *ϵ* = 0.01, *R* = 110, *m* = 10, *b* = 8, *l* = 10, *a* = 8. (b) Like (a) with *P*(*E*(*ρ*)) calculated using simulations instead of the Sanov bound and where the total number of reads *R*_*all*_ = 1000.

We note that the bound achieved by using Sanov’s theorem is not tight and therefore present an alternative approach for finding *ρ* using empirical simulations. Fig. 6b presents the probability (1 − *P*(*E*(*ρ*))) calculated like in Fig. 5 by 100,000 instances of simulating the multinomial distribution with *R*_*all*_ = 1000. Clearly, this method yields a tighter bound on the decoding probability while also requires analyzing less read overall.

### D. A tool for determining the required sequencing coverage

We have developed a tool designed to calculate the necessary sequencing coverage for DNA-based data storage systems.

#### Parameters, Input, and Output

The tool gets as parameters the sequence design and coding schemes and computes the required sequencing coverage for a desired confidence level. Specifically:

Design parameters:

- *K* – Total number of unique k-mers in each position.
- *t* – The required threshold on the number of observed occurrences of each of the k-mer
- *m* – sequence length
- *b* – the number of letters required to be successfully decoded in each sequence
- *l* – The number of sequences in the message
- a – The number of sequences required to be successfully decoded
- *ϵ* – Error probability of observing an invalid k-mer Input:
- *δ* – acceptable failure rate

Output:

- *R*_*all*_ – required sequencing coverage

#### Description of tool run

Fig. 7 presents a high-level description of the tool workflow. Given the design parameters *K, t, ε, m, b, l*, and *a*, the tool finds a threshold *ρ* and a total number of reads *R*_*all*_ for which conditions (a) and (b) hold for the input confidence level 1 − *δ* (Part C). First, *ρ* is found such the decoding of at least *a* sequences is ensured, 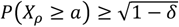 (Part C). This calculation requires the probability to decode a single sequence, *P*_*single*_ (*ρ, m, b*) (Part B) which uses the reconstruction probability of a single combinatorial letter *π*(*K, t*)(*ρ*) (Part A). Once *ρ* is determined, the algorithm searches for the required number of overall reads *R*_*all*_ that ensures 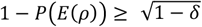 . This can be achieved using either using the bound from Sanov’s Theorem or using the empirical estimation of *P*(*E*(*ρ*)). When a value for *R*_*all*_ that satisfies the condition is found then the tool run exits outputting *R*_*all*_ to the user

**Fig. 7.**
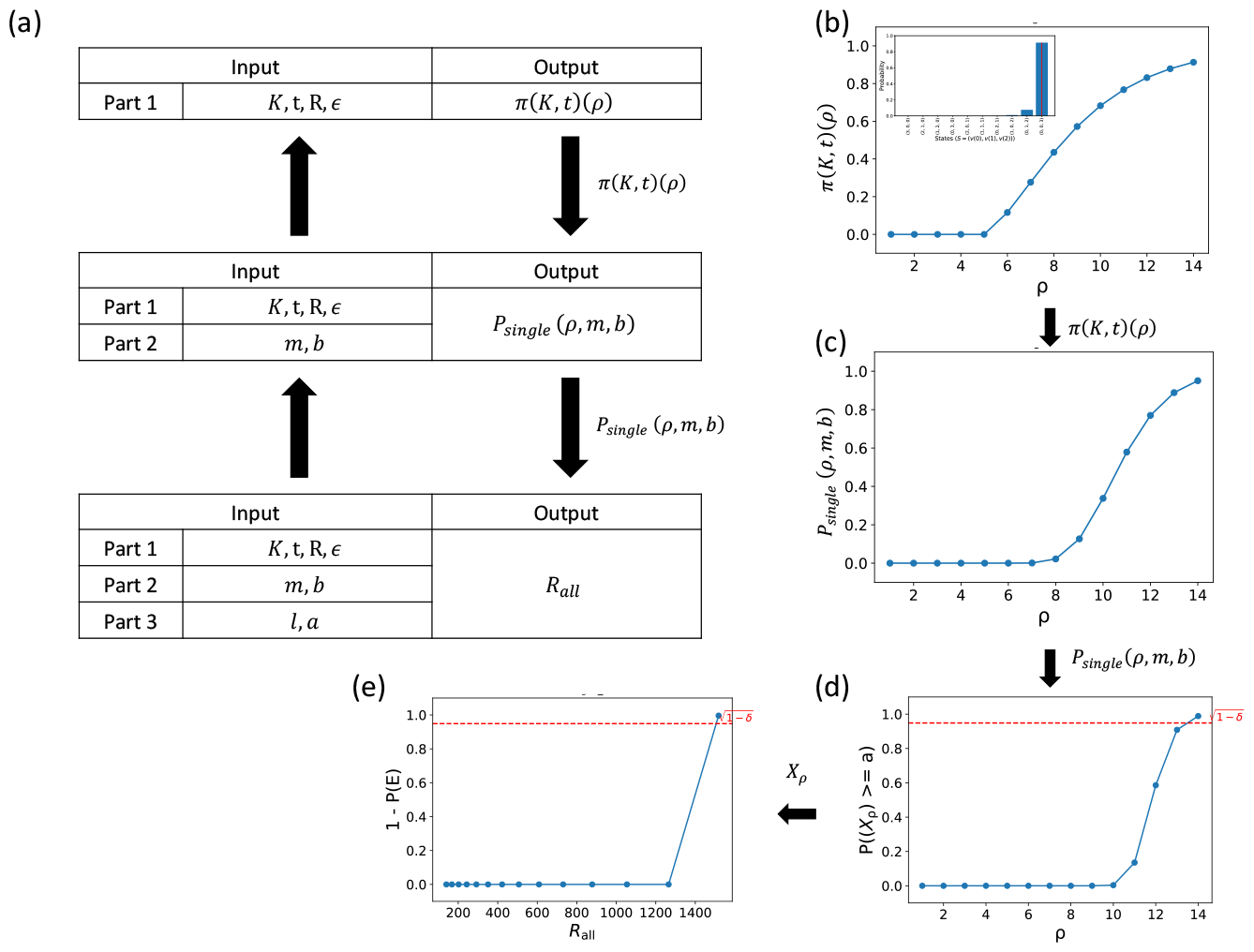
Complexity calculation tool workflow. (a) Overview of the tool’s run including internal dependencies, input parameters and outputs for each part. (b) Reconstruction probabilities of a single combinatorial position, *π*(*K, t*)(*ρ*), calculated using the coupon collector’s model (inset, like in Fig. 1) as a function of the threshold *ρ*. (c) Decoding probability for a full-length combinatorial sequence, *P*_*single*_ (*ρ, m, b*), calculated using the binomial model with the probabilities from (a) as input. Plotted as function of the threshold *ρ*. (d) Finding *ρ*. Full message decoding probability, *P*(*X*_*ρ*_ ≥ *a*), calculated using the binomial model for *X*_*ρ*_ obtained from (c). Plotted as function of the threshold *ρ*. The target confidence level 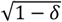 is presented in the dotted red line. (e) Finding *R*_*all*_ given the selected *ρ*. The probability of considering enough read distributions (across the *l* sequences), 1 − *P*(*E*(*ρ*)), based on either theoretical bound or the empirical calculation. Plotted as a function of *R*_*all*_ . The target confidence level 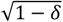 is presented in the dotted red line.

#### Example runs

To demonstrate the tool’s functionality, we used it to determine the required sequencing coverage for different sets of design parameters, similar to those used in [15], and for various confidence levels. These results are presented in Table 1. Clearly, increasing the desired confidence level (smaller values for *δ*) requires increasing the sequencing converge. Scaling up the system’s capacity by taking *l* to be 10 times larger results in a proportional increase in the *R*_*all*_ . Increasing the redundancy level (lower value for *a*) reduces the number of required reads to analyzed. We note that the different design parameters influence both the threshold *ρ* and the sequencing coverage *R*_*all*_ . While *R*_*all*_ is affected by all the design parameters, *ρ* is primarily affected by *m* and *b*. These findings underscore the importance of carefully selecting system parameters to optimize the efficiency and reliability of DNA-based data storage systems. Future work may explore the boundaries of these parameters to further enhance system performance.

**Table 1.**
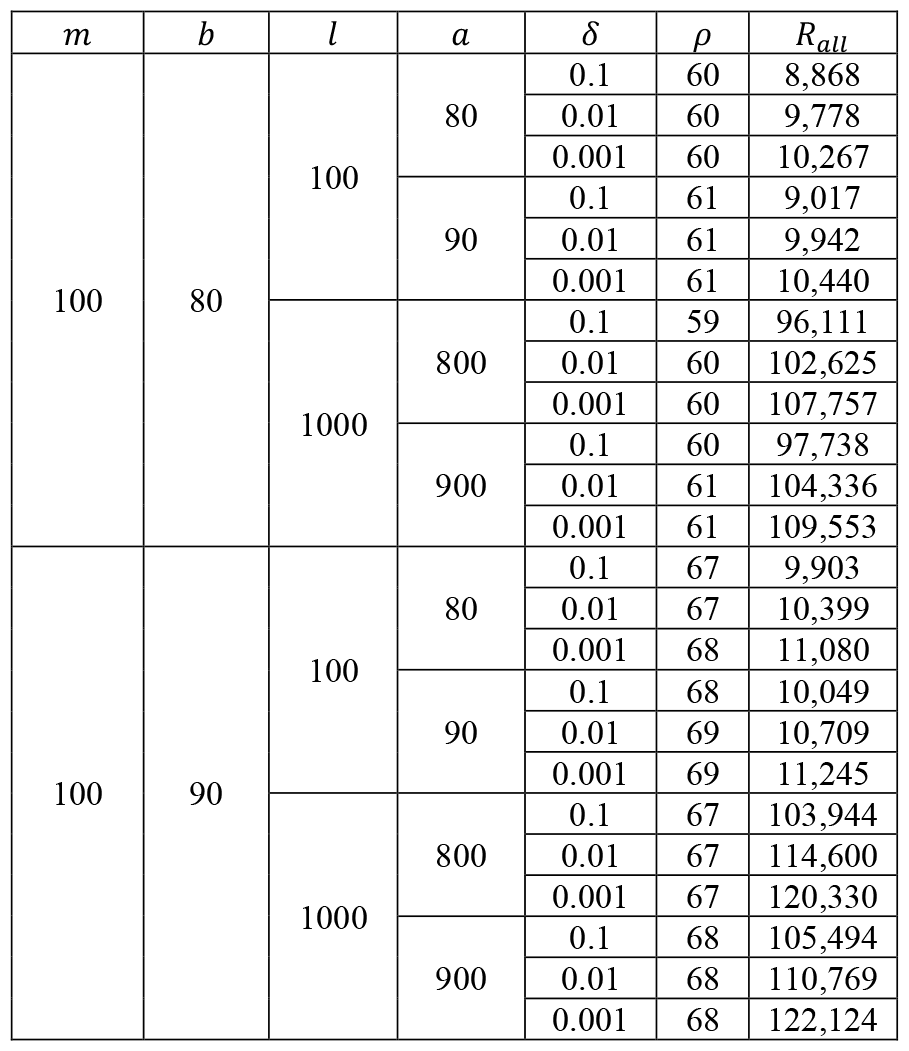
Required sequencing coverage for different design parameters and confidence levels.

## IV. Discussion

Our study presents a novel model for analyzing coverage depth in DNA-based data storage, particularly focusing on combinatorial DNA encoding. We use the coupon collector’s problem framework to model the reconstruction of combinatorial letters from sequencing data. We present a Markov Chain (MC) formulation for the calculation of the decoding probability and provide a tool for computing the probability. This solution is, however, limited in its scale due to the size of states space. Further work may be done to allow scaling up this model, either by developing more efficient computations or by developing approximation to the model.

One of the key aspects of the combinatorial approach is the strategic selection of Ω to consist easily distinguishable k-mers. This, together with the use of a threshold *t* > 1 in the reconstruction algorithm (See Algorithm 1) effectively mitigate k-mer mixup errors, as was demonstrated in [15]. We therefore chose to ignore k-mer mixup error in the model used for the reconstruction probability.

We also present a unified model for analyzing coverage depth of a complete combinatorial storage system considering an inner-outer error correction model. We present theoretical bounds on the decoding probability using Sanov’s Theorem on the multinomial model for read distribution or using an empirical estimation.

We also provide a python tool for determining the sequencing depth required to achieve a desired confidence level for a system given design and encoding scheme. We demonstrate the tool’s results on a selection of design parameter sets.

Future exploration in DNA data storage will significantly benefit from further understanding and optimizing coverage depth and from further improving efficient combinatorial coding. These elements are key to enhancing data storage capacity and reliability, promising exciting advancements in the field.

## Appendix

### A. Evolution of Probability in the Coupon Collector Problem

The coupon collector parameters that are showed in the video are: *K* = 5, *t* = 2, *R* = 30.

A. Evolution of Probability in the Coupon Collector Problem Video K=5, t=2, R=30.gif

### B. Classical coupon collector problem

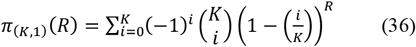

*π*(*K, t*)(*R*) is the probability of collecting all *n* unique coupons within *R* trials.

We will show that

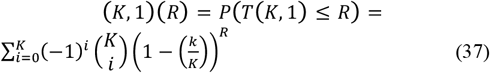

for the coupon collector’s problem, we can approach it using the principle of inclusion-exclusion. The formula calculates the probability of collecting all *n* unique coupons within *R* trials.

Let *A*_*i*_ be the event that the *i*-th coupon is not collected in *R* trials.

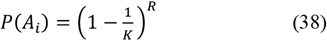

Let 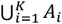 be the probability of not collecting at least one coupon in *R* trials.

Note that we are interested in:

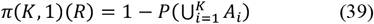

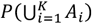 is calculated using the principle of inclusion-exclusion.

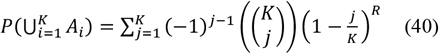

And finally,

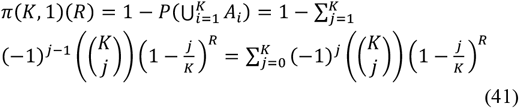

This follows from:

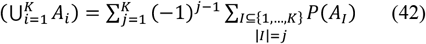

Where *A*_*I*_ = ⋂_*i*∈*I*_ *A*_*i*_

For *j* = 1:

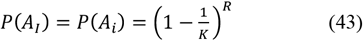

For *j* = 2:

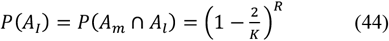

And generally

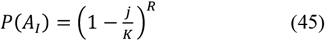

And clearly:

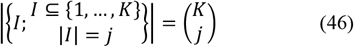

### C. Proof of the Lemma for the Sanov bound

Lemma:

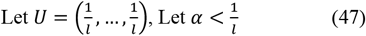

Let 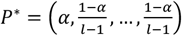, then

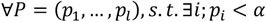

We have

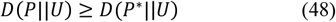

Proof:

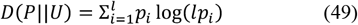

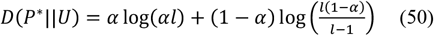

We solve:

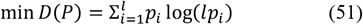

Subject to:

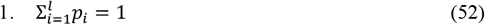

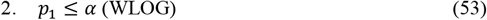

Therefore, the Lagrangian is:

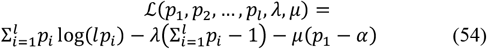

The KKT conditions are:

1. Stationarity 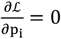 :

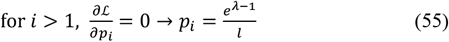

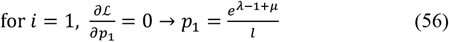
2. Primal feasibility:

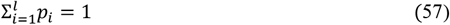

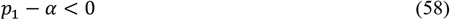
3. Dual feasibility:

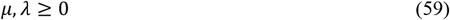
4. Complementary slackness:

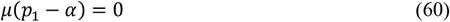

Expressing λ using the primal feasibility (57):

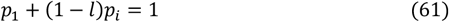

Substituting *p*_*i*_ and *p*_1_ from (55)(56):

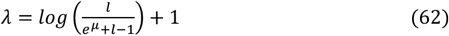

Expressing *p*_1_ with λ from (62):

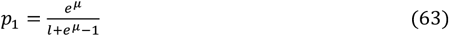

Expressing *μ*:

1. If *p*_1_ ≠ *α*, then *μ* = 0.
2. If *p*_1_ = *α*, then *μ* can be non zero.

If *p*_1_ = *α*, we substitute *α* for *p*_1_ in (63):

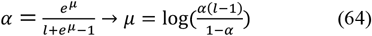

Using the original expressions for *p*_*i*_ from (55), and substituting *μ* we expressed in (64), we get:

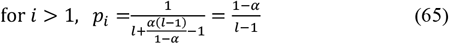

and recall that *p*_1_ = *α*

If *p*_1_ ≠ *α*, we substitute *μ* = 0 for *p*_1_ in (56):

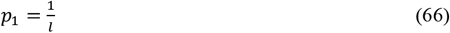

Using the original expressions for *p*_*i*_ from (55), and substituting *μ* = 0, we get:

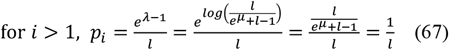

And we get the trivial solution *P*^*\**^ = *U* which does not satisfy the condition *p*_1_ < *α*.

Therefore, we proved that *D*(*P*|*U*) ≥ *D*(*P*^*\**^|*U*).

